# High-throughput transposon mutagenesis defines the essential genome of diverse phages

**DOI:** 10.64898/2025.12.19.695335

**Authors:** Natalie Kyte, Manuela Fuchs, Leah M. Smith, Peter C. Fineran

## Abstract

Phages are important drivers of bacterial evolution with therapeutic potential as antimicrobials. However, gaps in our understanding of phages and our inability to rapidly engineer them with new genetic cargo hinders progress towards phage-based therapies. To address the lack of unbiased, genome-wide mutational tools for phages, we developed transposon mutagenesis employing CRISPR-anti-CRISPR (Acr)-based selection and deep-sequencing (Phage Tn-seq). Transposon mutagenesis was effective for phages with unmodified or hypermodified genomes and a jumbo phage that protects its DNA within a nucleus. Phage Tn-seq enabled phage gene essentiality assignment consistent with structural proteomics and core gene conservation. Insertion biases allowed prediction of transcriptional direction and early injected phage DNA regions. We exploited the method to rapidly deliver new cargo to phage genomes in just a few days and used an AI-designed Acr to expand the phage transposon toolbox. Phage Tn-seq is a versatile tool to advance our understanding and applications of phages.

## Introduction

Bacteriophages (phages) are viruses that specifically target bacteria and play a crucial role in global biogeochemical cycles by driving bacterial turnover^1^. Their ability to infect and kill their specific host bacteria, even those resistant to antibiotics, has led to a resurgence in interest in their use as antimicrobials in humans (phage therapy), in food safety and in agriculture^2,3^. Despite recent high-profile human phage therapy successes^2,4^, many past interventions failed for unknown reasons, but likely due to insufficient knowledge of phage-host interactions. Indeed, phages are incredibly diverse and, except for few model phages, their biology remains poorly understood, with many of their genes lacking assigned functions. Moreover, the presence of extensive and often uncharacterised defence and anti-defence systems further complicates functional interpretation and reflects how much of phage–host biology remains unresolved.^5,6,7^. This gap in fundamental knowledge about phage genes and their genetic program limits the rational and synthetic biology application of phages as therapeutics and biotechnological tools. While traditional phage genome interrogation methods are largely low-throughput, genome-wide functional genomics approaches will facilitate a deeper understanding of phage biology, accelerating the resolution of gene function by enabling simultaneous, rather than sequential, analysis^8^.

Genome-wide silencing techniques have recently been developed to investigate gene essentiality in phages, distinguishing between genes essential or non-essential for phage replication under different conditions^9,10^. These methods include CRISPR-mediated gene knock-down (e.g. dCas12a and dCas13d-mediated CRISPRi-ART) and more recently, anti-sense oligonucleotides (ASOs)^11,12^. These approaches, although relatively effective for identifying essential genes via knock-down, can be costly, since they require the generation of guide RNAs/ASOs libraries to tile across a genome. These methods also rely on well-annotated phage genomes for the identification of suitable genomic contexts for gene silencing. Failure to fully silence genes can also result in incorrect assignment of essential genes as non-essential^9,11^.

The ability to mutate phage genomes is fundamental for assigning unknown gene functions or to introduce genetic ‘cargo’ to enhance phage functionality, such as to improve their therapeutic efficacy^13^. However, there are currently no genome-wide unbiased insertional mutational approaches for phages. Existing low-throughput phage engineering methods to generate single mutations rely on homologous recombination with template DNA, recombineering and CRISPR (clustered regularly interspaced short palindromic repeats)-Cas-based counter-selection^6–8^. Recently, PhageMaP was developed, which enabled CRISPR-directed gene essentiality mapping across phage genomes via homologous recombination^10^. Other approaches allow *in vitro* synthesis of engineered phages but are currently limited by genome size and restricted to a few model bacteria^8,14^. In addition, engineering phages with genetic cargo typically involves identifying ‘safe’ genomic sites to ensure phage function is not impaired. All these methods are time-consuming, costly, lack throughput and are biased by their directed nature.

We have developed a genome-wide phage random transposon mutagenesis and sequencing method to assess phage gene essentiality in a single experiment (Phage Tn-seq). This technique also enables rapid delivery of genetic cargo to phages without prior knowledge of ‘safe harbour’ sites. In bacteria, transposon insertion sequencing is a largely unbiased, cost-effective, and high-throughput approach to investigate gene essentiality or fitness. A saturated random transposon mutant library is generated, grown under one or more conditions and the locations of transposon insertions determined by deep-sequencing (e.g. Tn-seq, TraDIS)^15,16^. This allows the identification of non-essential genes with insertions and essential genes that remain undisrupted, which is a strategy we have previously applied to identify bacterial genes required for phage infection^17^.

Here, we developed Tn-seq for phages by leveraging anti-CRISPR (Acr) as a positive selectable marker in the presence of CRISPR-Cas counter-selection. The absence of suitable counter selection was previously a barrier for performing Tn-seq in phages, since the antibiotic selection methods used in bacterial Tn-seq are ineffective against phages. Using a Tn*5* transposon containing AcrVIA1 for phage mutagenesis, followed by Cas13a counter-selection and deep-sequencing of transposon junctions, we defined essential and non-essential genes in multiple phages in a non-model bacterium (*Serratia*/*Progidiosinella*). We achieved efficient phage genome mutagenesis of modified and hypermodified double-stranded DNA (dsDNA) phages and a nucleus-forming jumbo phage^18^. We further demonstrate the broader utility of the Tn*5* phage transposon as a rapid platform for cargo loading. As a proof of principle, we generated mEGFP expressing phages, which were visualised by confocal microscopy. We also show alternative cargo loading of an artificial anti-CRISPR, enabling enrichment of transposon insertions in hypermodified *Prodigiosinella* and *Escherichia coli* phages, demonstrating efficient transposition across distinct phages and host genera. Phage Tn-seq is a powerful method for high-throughput functional genomics and the exploration of gene essentiality in diverse phages.

## Results

### Development of an anti-CRISPR transposon and Cas13 counter-selection strategy

Our goal was to generate a phage transposon mutagenesis and counter-selection strategy that was applicable to diverse phages. The procedure involves mutagenesis of a phage upon infection of a host bacterium harboring a plasmid with a transposon encoding an anti-CRISPR protein (**Fig. 1A**). The resulting phages consist of a mixture of transposon mutants and parental phages. Mutants can then be enriched by growth on a counter-selection strain expressing a CRISPR-Cas system targeting the phage of interest. Only phages harboring a transposon and expressing an anti-CRISPR will survive. The locations of the transposons in the phage genomes can be identified through deep-sequencing, allowing determination of non-essential and essential phage genes.

**Figure 1.**
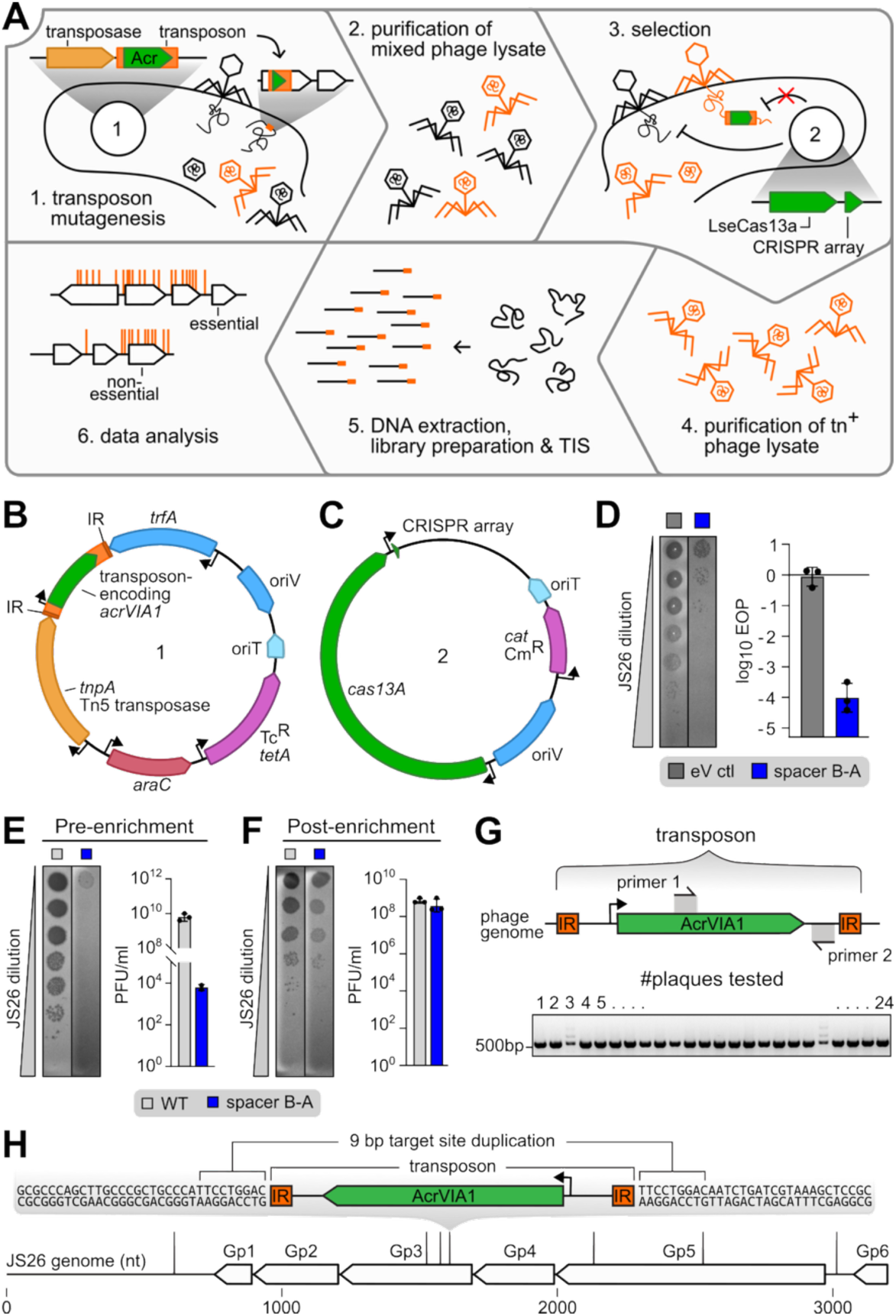
An anti-CRISPR transposon and Cas13a counter-selection generates transposon mutants in a dsDNA phage genome. **(A)** Schematic of the phage transposon mutagenesis. (1) Bacteria harboring an Acr transposon (plasmid #1, **B**) are infected with the phage of interest and phage production results in a mixture of transposon mutant and un-mutagenised phages that are (2) harvested. (3) The mixture of un-mutagenised phages and Tn^+^ mutant phages are then used to infect bacteria carrying a Cas13a counter-selection plasmid (plasmid #2, **C**) expressing gRNA(s) against the target phage. (4) The resulting enriched Tn^+^ mutant phages are purified, (5) DNA is extracted and libraries prepared and sequenced, and (6) sequencing data analysed to determine essential and non-essential genes/genomic regions of the phages. (**B**) Map of the plasmid pPF4213 containing Tn*5*-AcrVIA1, including the transposase under arabinose-inducible (AraC) control, an origin of conjugative transfer (oriT), a broad host-range origin of replication (oriV) and selectable marker (Tc^R^). **(C)** Map of the Cas13a counter-selection plasmid pPF3929 with oriT and oriV as per (B) and a selectable marker (Cm^R^). **(D)** Efficiency of plating (EOP) of phage JS26 on *Prodigiosinella* carrying either an empty vector control (grey; pPF3929) or the anti-JS26 Cas13a counter-selection plasmid (blue; pPF4421). Data are the geometric mean of n=3 independent biological replicates ± geometric standard deviation (SD), and individual data points are shown. **(E & F)** Phage titers following Tn5A-AcrVIA1 transposon mutagenesis either **(E)** pre-or **(F)** post-enrichment with Cas13a counter-selection. A representative image of phage spot dilutions of n=3 independent biological replicates is shown respectively. Error bars represent the mean ± SD of n=2-3 technical replicates and individual data points are shown. **(G)** Schematic of PCR verification of transposon insertions (top) and PCR screening of Phage plaques (bottom) picked following plating of the transposon mutagenesis lysate on *Prodigiosinella* carrying the anti-JS26 Cas13a counter-selection plasmid (pPF4421). **(H)** Mapping of transposon insertion sites for individual phages determined by whole genome sequencing. The 9 bp target site duplication resulting from Tn*5* transposition is shown (top).

We opted for the Tn*5* transposon because it inserts in a largely unbiased manner, promoting the generation of high-density mutant libraries, the analysis of which is supported by well-established methods^18,19^. Since some phages evade DNA-targeting CRISPR via diverse mechanisms^20,21^, we chose to use *Listeria seeligeri* (*Lse*) Cas13a. Cas13 is a type VI CRISPR-Cas system that targets and cleaves RNA, thereby overcoming most CRISPR evasion strategies, including DNA modification and nucleus formation in jumbo phages^22,23^. Cas13a has been shown to block diverse *E. coli* phages and a *P. aeruginosa* jumbo phage, while its anti-CRISPR AcrVIA1 has been used as a selectable marker to generate defined mutations^22,24^.

To generate a widely applicable Acr-transposon platform (**Fig. 1B**), we created a transposon encoding AcrVIA1, driven by a strong constitutive promoter, on a broad-host range plasmid with an RK2 origin of replication that functions in diverse bacteria^25^. In addition, to allow efficient introduction to different bacteria, even those with low competence, an RP4 origin of transfer (oriT) for conjugative delivery was included. To minimise Tn*5* mutagenesis of the host bacterium prior to phage introduction, transposase expression was controlled by the tightly inducible *araBAD* promoter under AraC repression. For selection of transposon mutant phages, we generated a counter-selection plasmid for Cas13a expression containing a BsmBI cloning site for insertion of guide RNAs against selected phages, in a streamlined one tube golden gate reaction (**Fig. 1C and Fig. S1A**).

### Anti-CRISPR efficiently selects phage transposon mutants in a non-model system

To test phage transposon mutagenesis, we chose the siphovirus JS26 and its non-model host *Prodigiosinella confusarubida* (formerly *Serratia* sp. ATCC 39006), for which it is currently the sole known genus representative^21^. JS26 is a member of the *Dunedinvirus* genus and has a 64 kb unmodified dsDNA genome, making it a relatively simple initial candidate phage to test phage Tn-seq^26^. We created two Cas13a counter-selection plasmids to verify their activity, each with a gRNA targeting either of two different genes (gp81 an exonuclease and gp5 a hypothetical protein). In efficiency of plaquing assays (EOPs), JS26 was targeted by both spacers (spacer A & B) with a 10^3^-10^4^-fold reduction (**Fig. S1B**). To reduce the background of apparent escape mutant phages, we generated Cas13a plasmids encoding both gRNA in either order (spacer A-B and B-A) and repeated the EOP assays (**Fig. S1B**). Spacer B-A further decreased phage infection by ~10-fold and was chosen for subsequent counter-selection steps (**Fig. 1D**).

Having established the Tn*5*-AcrVIA1 transposon mutagenesis and verified the counter-selection, we tested mutagenesis of JS26. Bacteria with the Tn*5*-Acr plasmid were pre-induced for 2 h to express the transposase followed by phage infection in liquid to facilitate Tn*5*-Acr transposon insertion into the phage genome. Phages were harvested and used to infect bacteria containing the Cas13a counter-selection plasmid, resulting in enrichment of phages containing a transposon insertion. By plating the mutagenised phages pre- and post-enrichment on Cas13a counter-selection, a clear enrichment in Acr containing phages (i.e. Cas13a-insensitive) was observed (**Fig. 1E,F**). To confirm that phages had acquired the Tn*5*-AcrVIA1 transposon, we picked individual plaques from Cas13a counter-selection plates and verified the successful transposon insertion via PCR (**Fig. 1G and Fig. S1C**). Finally, we performed whole genome sequencing on individual phages, which revealed single Tn*5*-AcrVIA1 insertions in each phage genome in different genomic locations with the distinctive 9 bp target site duplication caused by the Tn*5* transposition mechanism (**Fig. 1H, Fig. S1D**)). In summary, we could show that anti-CRISPR transposon and Cas13 counter-selection can be used to generate multiple distinct Tn*5* mutations in the genome of an dsDNA phage.

### Phage Tn-seq facilitates a genome-wide view of gene essentiality

Having established that transposon mutagenesis of the JS26 genome was achievable, the triplicate post-enrichment transposon phage lysates (i.e. after outgrowth on the Cas13 counter-selection strain) (**Fig. 1F**), were subjected to DNA extraction, transposon junction Illumina amplicon library preparation and deep sequencing of Tn*5*-AcrVIA1 insertion sites (**Table S1**)^18^. Following enrichment, each replicate contained >330,000 reads and ~1400 to 2500 unique insertions per phage genome with an average of one unique transposon insertion per 32 nt. Combining all replicates resulted in 3765 unique insertions with one on average every 17 nt, demonstrating a high-resolution mapping of non-essential insertion locations (**Fig. 2A and Table S1)**. Of the 84 coding sequences in the JS26 genome, 47 (56%) were predicted as essential via the TraDIS pipeline (**Extended Data Table 1**)^19^. Apart from three genes (gp10, gp15 and gp40), most essential genes were located together in clusters (gp28-gp30, gp32-gp33 and gp46-gp84) (**Fig. 2A,B**). To compare these insertion distributions to the mutant pool prior to enrichment, we also sequenced phages after transposon mutagenesis but prior to outgrowth and counter-selection. Prior to enrichment, a greater proportion of reads was found in genes later classed as essential following Cas13 counter-selection, as some of these mutants likely failed to infect or replicate following initial mutagenesis and burst (**Fig. S2A-B**).

**Figure 2.**
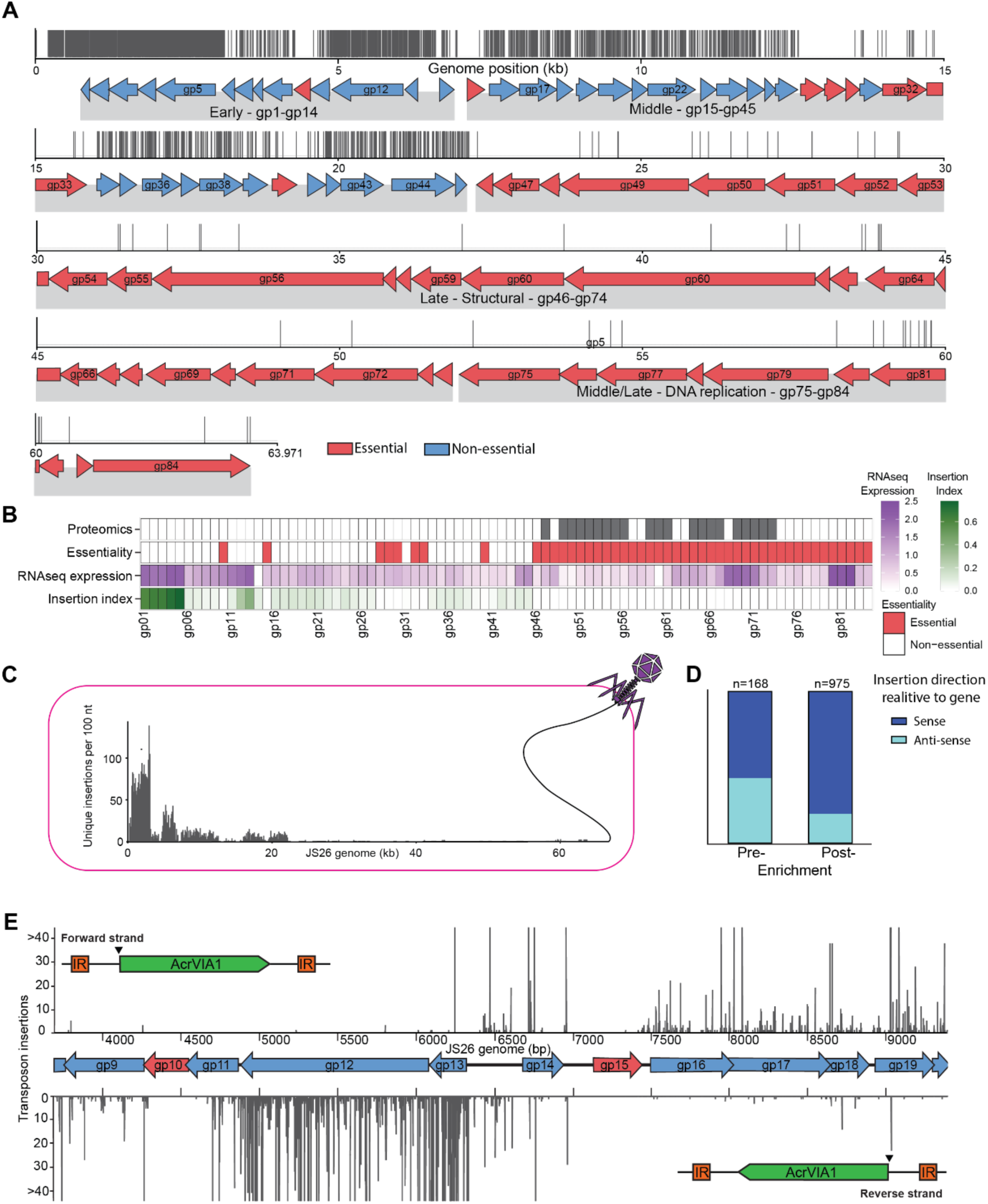
Phage Tn-seq provides a genome-wide map of JS26 gene essentiality. (**A**) Unique transposon insertions across the JS26 genome. Essential genes are shown in red and non-essential in blue. Early, middle, late, and middle-late gene clusters are indicated. (**B**) Summary of transposon insertion index, gene essentiality, gene expression and structural proteomics in JS26. See **Extended Data Table 1.** (**C**) Over-represented insertion region at the first injected portion of the JS26 genome shown relative to the predicted order of genome entry. Unique insertions are binned in 100 nt bins. (**D**) Number of unique insertions in genes relative to gene orientation, except the hot-spot region (i.e. the transposon is either sense or anti-sense to gene transcription; n= the total number of unique insertions within genes). (**Fig. S2E** shows data including the insertion hotspot). (**E**) Total transposon insertions in a representative region of the JS26 genome. Insertions in the forward orientation are shown above and insertions in the reverse orientation below. Essential and non-essential genes are evident, even within operons (e.g. gp10).

To validate the predicted essential genes, we first compared the essentiality data to the genome annotation, RNA-sequencing infection time-course data and proteomic data of the mature structural proteins in the JS26 virions (**Fig. 2B, Extended Data Table 1**)^26^. JS26 encodes an operon containing 28 predicted structural proteins (transcribed from gp73 to gp46), most (75%) of which were detected in the mature JS26 virions. Importantly, all proteins detected in the mature virions were predicted to be essential, corroborating the ability of phage Tn-seq to identify essential genes. Seven structural proteins (gp46, gp48, gp57, gp58, gp62, gp63 and gp68) were also deemed essential but not detected in mature virions. Gp46 is a predicted Rz-like spanin involved in cell lysis for phage release^27^, Gp57 and Gp58 are predicted tail assembly chaperones and Gp62 and Gp63 are predicted pre-tape measure and tail length tape measure proteins, respectively. Gp48 and Gp68 have no predicted function; however, Gp68 is conserved across several phages, suggesting it may be essential in other genera. Another cluster of genes (gp74 to gp84) was also classified as essential, which correlates with their organisation as two middle-late expressed operons involved in DNA replication (**Fig. 2B**). In agreement with the essentiality predictions, dCas9 CRISPRi knockdown of one of these genes (gp81) leads to decreased phage fitness^26^.

Most early expressed genes (12 of the 13) were classified as non-essential, except for gp10. Since many early-expressed phage genes are predicted to have roles in anti-defence or host takeover^7^, we hypothesised that JS26 Gp10 may function similarly. Within the middle-expressed cluster, eight of 30 genes were essential. Of these, five essential genes were clustered together (gp28-gp30 and gp32-gp33) and two additional hypothetical genes (gp15 and gp40) were separate and flanked by non-essential genes. The gp28-gp33 region is predicted to encode proteins involved in DNA transcriptional regulation, DNA modification and potentially DNA repair. Gp30 is a predicted winged helix-turn-helix (wHTH) transcriptional regulator, Gp32 is related to Dcm methyltransferases and Gp33 is an exonuclease recombination-associated protein. Gp28 and Gp29 have no predicted function. Overall, phage Tn-seq is an efficient genome-wide method to identify essential and non-essential regions of phage genomes.

### Transposon strand bias and insertion frequency reveal transcriptional direction and early injected phage DNA

To investigate if the transposon insertion orientation or frequency provided further insight into the biology of phage genome organisation, we correlated insertion abundancies with gene expression patterns. We observed highly abundant and unique insertions in the early, highly expressed region of JS26 both with or without Cas13 counter-selection (**Fig. 2A,C and Fig. S2A**). In bacterial Tn-seq, Tn*5* insertion is not biased by gene expression levels^15^. Indeed, the insertion indices for the remainder of genes (middle and late) before Cas13 counter-selection did not correlate with expression (RNA) levels (**Fig. S2C**), indicating another factor leading to increased transposition in the early region (gp01 to gp13). Phage JS26 belongs to the *Casjensviridae* family, members of which use cos-site DNA packaging^28,29^. Indeed, sequence analysis of the JS26 genome ends from mature virions revealed defined ends starting before gp01 and ending after gp84, consistent with cos-type packaging (**Fig. S2D**). Phage lambda is the model cos-type phage and has been shown to inject the same initial part of the genome from each virion via fast self-repulsion from the capsid^30^. A slower second step involves anomalous diffusion of the remainder of the phage genome into the bacterial cytoplasm^31,32^. This mechanism is consistent with the highly mutagenised region of JS26 at the start of the genome, which in theory is exposed to Tn*5* insertion for a longer time period (**Fig. 2C**). These genes are small, non-essential and expressed highly within 5 mins of phage infection (**Fig. 2B**), consistent with early injection and likely provide anti-defense or host-manipulation functions.

Next, we analysed the orientation of transposon insertions. Following outgrowth and Cas13 counter-selection, the insertion direction of Tn*5*-AcrVIA1 in non-essential genes was almost completely correlated with gene directionality (**Fig. 2D,E and Fig. S2E)**^26^. In contrast, this bias was not observed for genes prior to counter-selection by Cas13. Therefore, transcription from native phage promoters is likely driving Acr expression in the transposon and allowing phage survival in the presence of Cas13 counter-selection. As such, investigating transposon insertion directionality could be useful in poorly characterised phages to help assign gene orientation and the direction of transcription.

Transposons can cause polar effects on transcription, which may influence the expression of downstream genes in operons – a possible issue for determining gene essentiality in phages with a modular organisation like JS26. Therefore, we examined whether Tn*5*-AcrVIA1 could discriminate between essential and non-essential genes in an operon (**Fig. 2E**). We identified examples of operons where upstream genes were non-essential (e.g. gp11), and hence contained insertions, whereas downstream genes lacked insertions and were essential (e.g. gp10). This demonstrates that phage Tn-seq with Tn*5*-AcrVIA1 allows read-through transcription and can distinguish essential from non-essential genes in operons. In summary, the insertion bias – both direction and frequency – of phage Tn-seq can uncover transcriptional direction, reveal early injected regions of phage genomes and essential and non-essential genes within operons.

### Phage Tn-seq functions with a nucleus-forming jumbo phage

Having demonstrated the efficiency of phage Tn-seq with JS26, we wanted to test it on a phage with a more complex lifecycle. Phage PCH45 is a nucleus-forming jumbo phage of the *Chimalliviridae* family with a dsDNA genome of 212,807 bp^21^. Members of this family include the well-studied phiKZ and related phages of *Pseudomonas* species^33–35^. PCH45 and other *Chimalliviridae* form a proteinaceous nucleus-like structure during infection that houses DNA replication and transcription, and protects their genomes from DNA-targeting defenses, such as CRISPR-Cas^21,36,37^. The phage RNA exits the nucleus for translation and hence renders these phages vulnerable to RNA-targeting CRISPR-Cas systems, including Cas13^21,36,38^. Prior to nucleus formation, *Chimalliviridae* produce an early phage infection (EPI) vesicle that is membrane bound and transcriptionally active^39^. Due to the protected nature of PCH45 DNA within an EPI and the nucleus, it was unclear whether phage Tn-seq would be effective for a nucleus-forming jumbo phage.

First, we established Cas13 counter-selection plasmids with gRNAs targeting different PCH45 genes and a dual gRNA plasmid was selected for subsequent experiments due to its stronger targeting efficiency (**Fig. S3A**). Phage Tn-seq was performed on PCH45, and the resulting transposon mutants were enriched using Cas13-based phage targeting, verified by PCR (**Fig. S3B**) and isolated phages demonstrated to fully overcome Cas13 immunity (**Fig. S3C**). Four transposon mutant phages were sent for whole-genome sequencing, revealing a single Tn*5*-AcrVIA1 insertion in different locations in each phage (**Fig. S3D**). Next, we deep-sequenced the transposon junctions in the enriched PCH45 transposon mutant pools, which revealed between ~200-500 unique insertions per pool (884 unique insertions, one every 240 bp on average for the combined replicates), which was insufficient coverage for gene essentiality analysis (**Fig. S3E and Table S2**). This low efficiency could be due to the EPI vesicle and/or phage nucleus which may reduce the access of the Tn*5* transposase and transposon to PCH45 phage DNA. In summary, Tn*5*-AcrVIA1 can access the jumbo phage DNA, suggesting that the barrier is not absolute and that the mutagenesis or sequencing could be scaled accordingly to obtain sufficient mutants.

### Transposase import into the jumbo phage nucleus improves Tn-seq efficiency

To improve the frequency of the transposon insertion into the jumbo phage genome, we created a fusion of PCH45 UvsX (Gp72), a RecA homologue, to the Tn*5* transposase protein (TnpA) (**Fig. 3A**). UvsX homologues localise into the phage nucleus during jumbo phage infection and can be used to deliver other genes into the nucleus when translationally fused^40,41^. First, since UvsX of PCH45 has not been studied, we created a UvsX-mCherry2 fusion and confirmed by confocal microscopy that it localises to the phage nucleus (**Fig. S3F**). Next, we tagged TnpA with mCherry2 and demonstrated its predominantly cytoplasmic distribution with partial nuclear localisation in PCH45 infected cells (**Fig. 3B**). Importantly, fusion of UvsX to mCherry2 and TnpA resulted in localisation inside the phage nucleus, indicating that the fusion mediated transposase import successfully (**Fig. 3B,C**).

**Figure 3.**
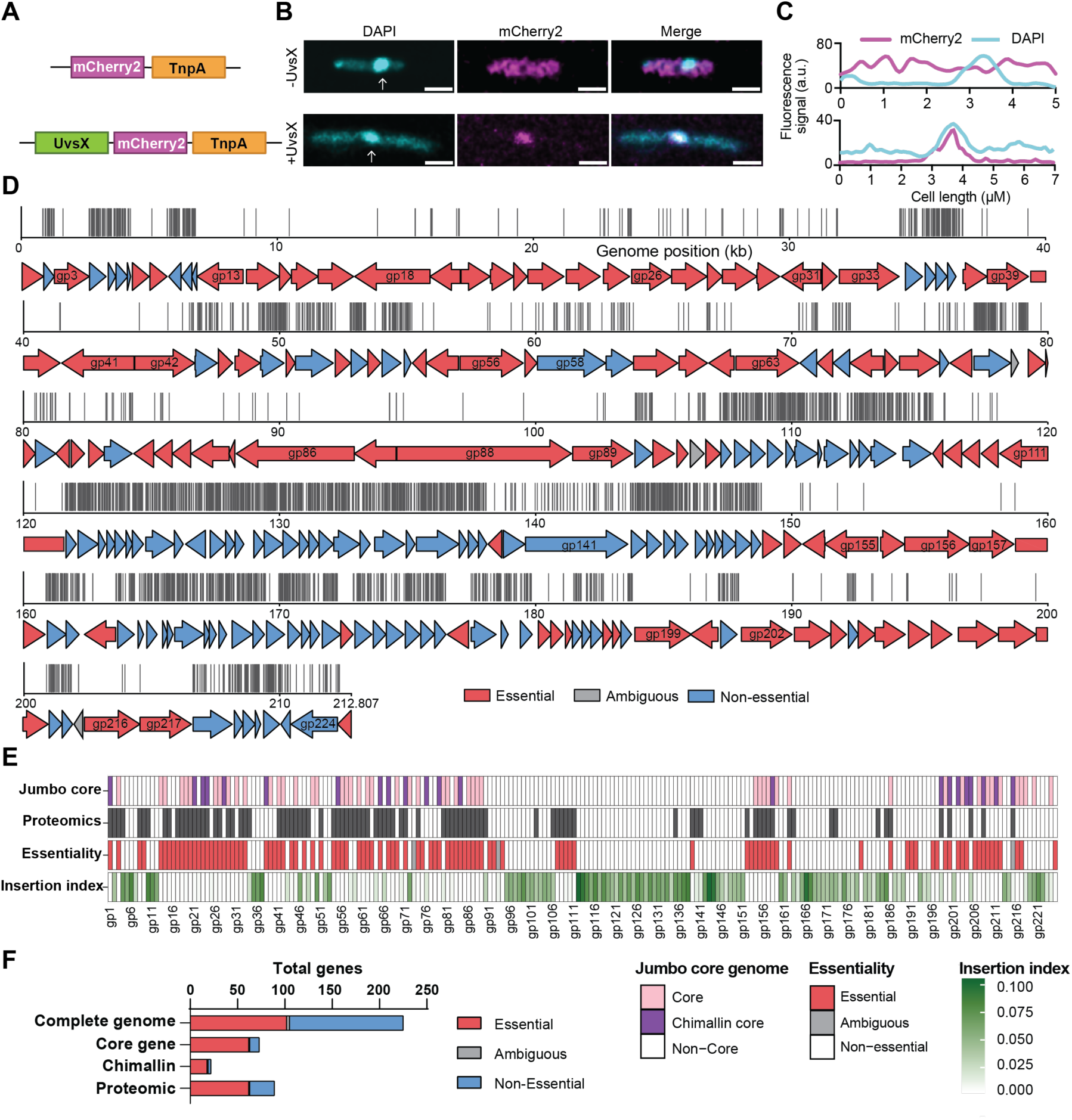
Phage Tn-seq provides a genome-wide essentiality map of the nucleus-forming jumbo phage PCH45. (**A**) Schematic of Tn*5* transposase (TnpA)-mCherry fusions with and without UvsX. (**B**) Confocal microscopy of *Prodigiosinella* harbouring a plasmid expressing a fusion of mCherry2-TnpA (pPF4355) or UvsX-mCherry2-TnpA (pPF4356) in the presence of PCH45 phage infection. DNA (cyan) was stained with DAPI and the protein fusions were tagged with mCherry2 (magenta). The PCH45 phage nucleus is indicated with white arrows. Scale bars are equal to 2 µM. (**C**) Fluorescence intensity plots show the distributions of the mCherry2 fusions (magenta) and DNA (DAPI, cyan) across the length of single cells. (**D**) Unique transposon insertions across the PCH45 genome. Essential genes are shown in purple, non-essential in blue, and ambiguous in grey. (**E**) Summary of transposon insertion index, gene essentiality, structural proteomics, and core genome predictions in PCH45. See **Extended Data Table 2 and Extended Data Table 3.** (**F**) Gene essentiality assignment to the complete genome, core genome predictions, nucleus forming genome predictions and proteomics.

Having demonstrated the UvsX-mediated transposase import into the phage nucleus, we tested a UvsX-TnpA fusion with the Tn*5*-AcrVIA1 transposon in Tn-seq against PCH45 (**Table S2**). Transposon insertions in PCH45 were verified via PCR and an improved mutagenesis using the *uvsX* fusion was observed by comparing titres of the phage mutant libraries generated using either the WT or UvsX-fused transposase on Cas13 counter-selection plates (**Fig. S3G**). Next, the mutant libraries were processed and analysed following deep sequencing of Tn*5*-AcrVIA1 insertion sites. Combining all UvsX-TnpA-generated transposon mutant library replicates, identified 3663 unique insertions in the PCH45 genome with one on average every 58 nt, demonstrating a high-resolution mapping of this jumbo phage. For further analysis we combined the 6 replicates of the UvsX-TnpA and the TnpA mutagenesis, increasing the total number of unique insertions to 3799 with an insertion every 56 nt (**Fig. 3D**). Of the 225 genes in PCH45, 102 (45%) were classified as essential, while 3 were ambiguous **(Fig. 3D,E, Extended Data Table 2)**. Therefore, at least 120 (53%) genes are likely non-essential, indicating that phages with large genomes carry considerable accessory genetic material. Transposon insertions were more widely distributed throughout PCH45 when compared with the JS26 distribution, which likely reflects less transcriptional organisation of *Chimalliviridae* genomes^42,43^. Similar to the Tn-seq of JS26, we observed a strong relationship between transposon insertion orientation with gene directionality (**Fig. S3H)**. In conclusion, phage Tn-seq can be used to predict essential and non-essential regions of nucleus forming jumbo phage genomes.

### PCH45 Tn-seq coupled with proteomics reveals essential and non-essential virion-associated proteins

We hypothesised that essential PCH45 genes predicted by Tn-seq should be overrepresented in those encoding virion proteins as these are likely essential structural proteins required for phage infection. To test this, we purified PCH45 particles and analysed their proteins by mass spectrometry. We identified 90 PCH45 proteins with at least two unique peptides, indicating their potential presence in mature phage virions (**Fig. 3E and Extended Data Table 3**). Of these proteins, 63 (70%) were predicted essential by phage Tn-seq. Therefore, proteins present in phage particles are more likely to be essential when compared with the total phage proteome (>70% *vs.* 45%) (**Fig. 3E,F**). As expected, many PCH45 virion proteins predicted as essential had important roles in phage viability. For example, proteomics and Tn-seq identified proteins involved in formation of the head (i.e., major capsid protein (gp33), internal head vertex proteins (gp23 and gp24), head maturation protease (gp81) and virion-associated head protein (gp162) and tail (i.e., tail tube (gp154), tail sheath (gp155), tail fibres (gp56 and gp60) and tail tip (gp19)). Similarly, PCH45 gp111 is an essential tail fibre protein and a homologue of ΦPA3 gp164 (a *Chimalliviridae*), which localises to the phage “bouquet” of assembled particles during phage maturation in the cell^44^. In addition, there was evidence of injected essential proteins, such as virion-associated RNA polymerase (vRNAP) subunits (gp3, gp87 and gp88). Interestingly, twenty six proteomic hits were not essential. Most have unknown functions or were less abundant, except gp58 and gp141, which are predicted tail fibre and tail proteins, suggesting they are dispensable under the conditions of the Tn-seq screen. Overall, proteomic data of PCH45 virions supports the Tn-seq essentiality predictions of phage structural or essential injected proteins.

### Genetic essentiality in PCH45 correlates with conservation across jumbo phages

We predicted that genes essential for PCH45 should be conserved across jumbo phages. A bioinformatic study compared jumbo phages of the *Chimalliviridae* family, and identified two overlapping sets of core genes (designated cg1-cg68); 1) those conserved across all jumbo phages “*core*” and 2) a sub-set of those conserved only in *Chimalliviridae* (“*chimallin-encoding core*”)^45^. Therefore, we compared the PCH45 essential genes (as determined by Tn-seq) with both conserved gene sets. There was a clear over-representation of both “*core*” and “*chimallin-encoding core*” genes within the essential genes. Indeed, ~86% of genes that were core to *Chimalliviridae*, with homologues in PCH45, were predicted as essential (**Fig. 3E,F**). Of the 21 *“chimallin-encoding core”* genes, 22 homologues were detected in PCH45, 18 of which were essential and one was ambiguous. Fifteen of the *“chimallin-encoding core”* genes were detected by proteomics (**Fig. 3E and Extended Data Table 3**). Other core genes were essential, but as expected not part of the virion due to their role during phage replication within the bacterial host. These include the proteinaceous shell of the phage nucleus ChmA (gp202; cg4), RNA binding protein ChmC (gp209; cg10), RNA polymerase beta subunit (gp203; cg5), nuclear shell import proteins PicA/Imp1 (gp211; cg12) and Imp3 (gp204; cg6) and the large terminase (gp159; cg66)^37,41,46,47^. Although not assigned within the “core” genome, the small terminase (gp77) and portal protein (gp206) were also essential. These results are consistent with conserved jumbo phage genes functioning in critical steps in the viral lifecycle and reinforces the efficacy of Tn-seq in essential gene identification.

This comparison also allowed us to identify non-essential core genes. Within the *“chimallin-encoding core”* genes, two non-essential (gp76 and gp205) and one ambiguous (gp215) gene have no known function, whereas the non-essential gp198 gene encodes an SH3 family protein. Within the entire set of “*core*” genes across all jumbo phages, seven were non-essential in PCH45. In some cases, there are multiple paralogues, with one being non-essential. For example, gp56, gp57 and gp58 are tail fibre proteins that are similar to cg42, with only gp58 being non-essential. Likewise, head maturation proteins gp81 and gp220 are related to cg60, with gp220 non-essential. A thymidylate kinase (gp50; cg32) is involved in DNA biosynthesis^48,49^ and is dispensable, potentially due to functional redundancy with a host-encoded thymidylate kinase. Although UvsX localises to the phage nucleus (**Fig. S3F**), it was non-essential, which is consistent with the dispensable nature of a related recombinase in T4^50^. Although not part of the core genome, the tubulin homologue PhuZ (gp187) was non-essential in PCH45, consistent with its dispensable role in phiKZ^24^.

We also identified 21 essential PCH45 proteins that are not members of the core jumbo phage genome. Gp9 has no predicted function but is a homologue of ΦPA3 gp078, which localises to the phage nucleus^44^. The essential gp107-gp111 operon was detected by proteomics and includes the predicted tail fibre (gp111) and tail assembly chaperones (gp107 and gp110). In addition, cell lysis cassette genes were essential, with gp152, a putative lytic transglycosylase, and gp54, which is predicted to encode an Rz lysis/spanin protein^51,52^. Taken together, PCH45 essential genes are enriched for conserved *Chimalliviridae* core genes, as well as virion proteins, while non-essential genes are frequently hypothetical or associated with functional redundancy. Overall, Tn-seq defined a core set of predicted genetically essential functions in PCH45.

### Phage transposon mutagenesis enables rapid loading of genetic cargo

In addition to the ability to define the essential genome of phages, phage transposon mutagenesis allows the rapid loading of genetic cargo into phage genomes without prior knowledge of ‘safe harbour’ sites. For example, we already demonstrated the generation of phages with broader resistance to CRISPR-Cas (Cas13) immunity through our Tn*5* delivery and in theory other genetic cargo could be delivered to phage genomes. This could prove useful for loading phages with other anti-defence genes, or genes that improve the therapeutic potential of phages. Furthermore, the ability to fluorescently label phages to follow their infection is useful for studying phage-bacterial dynamics at the single cell level. To test if we could rapidly deliver new cargo to a phage, without knowing a ‘safe’ insertion site, we generated a Tn*5*-mEGFP-AcrVIA1 transposon (**Fig. 4A, Fig. S4A**) and performed mutagenesis of PCH45. By using a streamlined mutagenesis procedure, only 2 days were required to identify phages harbouring Tn*5*-mEGFP-AcrVIA1 by PCR. By day 3, we confirmed with confocal microscopy that jumbo phage infected cells (discernable via nucleoid formation) were likewise positive for mEGFP (**Fig. 4B**). In parallel, we also mutagenised phage JS26 with Tn*5*-mEGFP-AcrVIA1, enabling streamlined identification of infected cells, which can otherwise be challenging for phages that do not produce easily identifiable hallmarks of infection (**Fig. S4B**). This also demonstrates that swapping *megfp* for other cargo will enable rapid engineering of phages to express any genetic cargo of potential therapeutic benefit.

**Figure 4.**
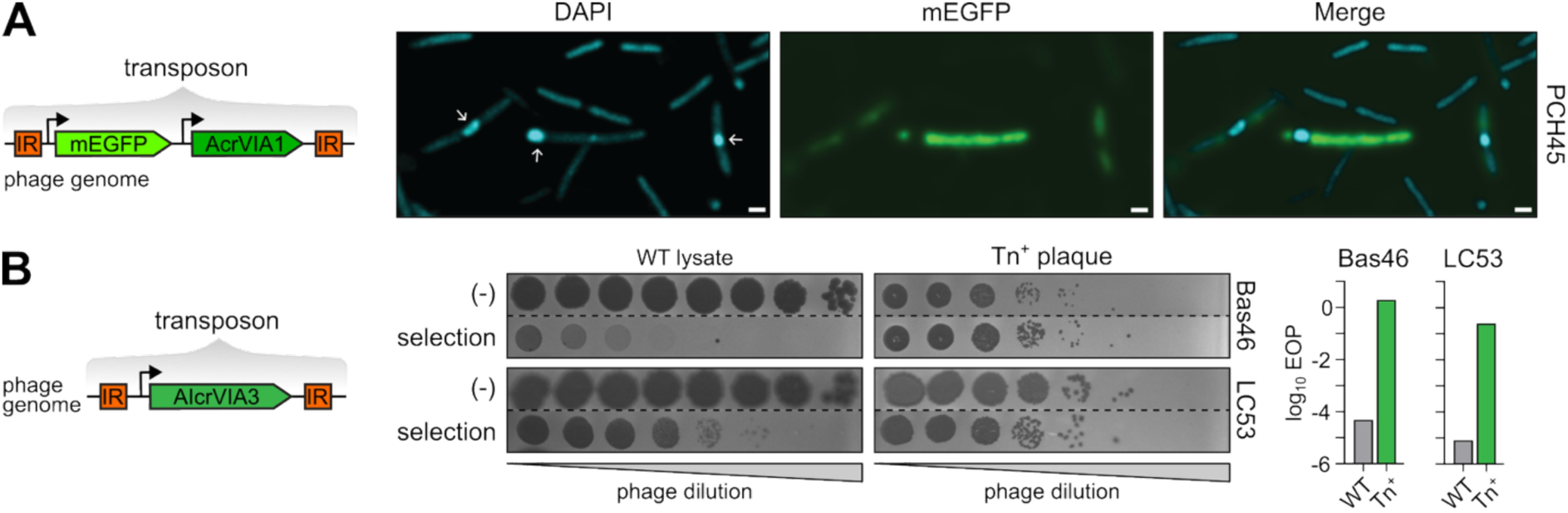
Modified transposons efficiently deliver cargo to generate phages with fluorescent labels or alternative anti-CRISPRs. (**A**) Map of the plasmid pPF4485 Tn5 containing mEGFP and AcrVIA1 (**B**) Confocal microscopy of *Prodigiosinella* infected with PCH45::Tn*5*-mEGFP-AcrVIA1. Phage DNA (DAPI) is shown in cyan and phage-expressed *mEGFP* (mEGFP) is shown in green. Infected cell jumbo phage nuclei are indicated with white arrows. Scale bars are equal to 1 µM. (**C**) Map of the plasmid pPF4881 Tn5 containing an AI designed Acr protein. (**D**) Phage spot dilutions of phage plaques picked following plating of the transposon mutagenesis lysate on *E. coli* DH10β carrying the anti-Bas46 LbuCas13a counter-selection plasmid (pPF3924), or on *Prodigiosinella* anti-LC53 LbuCas13a counter-selection plasmid (pPF3432 and pPF3434 mixed at a 50/50 ratio), phage lysate of the wildtype phage were used as controls. (**E**) Bar graphs comparing the EOP of WT lysate and a transposon mutant calculated as ratio of plaques on the phage selection strain relative to the number found on the wildtype stain control (-) for both Bas46 and LC53 respectively.

### An artificial Acr transposon and LbuCas13a enables mutagenesis of phages with hypermodified DNA in different genera

Phage mutagenesis would benefit from the ability to perform secondary rounds of mutagenesis to deliver additional genetic cargo or examine the effects of double mutations on fitness. However, once Tn*5*-AcrVIA1 has been delivered, alternative selection markers are required. Despite the many Cas13 variants, known Acrs are limited^53^. Recent advances in artificial intelligence have enabled the *de novo* design or discovery of anti-CRISPRs, including candidates against Cas13a and Cas13b^54,55^. Using a recently published AI-Acr (AIcrVIA3), we created a new transposon (Tn*5*-AIcrVIA3) that inhibits LbuCas13a (**Fig 4C, Fig. S4C**). Having demonstrated that phage Tn-seq works against phages with unmodified DNA (JS26) and DNA protected in a nucleus (PCH45), we now sought to test transposon mutagenesis in phages with hypermodified DNA. We performed mutagenesis on *Prodigiosinella* phage LC53 and *Escherichia coli* phage Bas46, which contain DNA with either 5-arabinose-hydroxy-cytosine or 5-arabinose-arabinose-hydroxy-cytosine, respectively^23^. Plaques were picked following mutagenesis and counter-selection on strains harbouring LbuCas13a and spacers targeting either phage, and transposon insertion confirmed by PCR (**Fig. S4D**). Next, we demonstrated that, by acquiring the AIcr, these phages had gained the ability to overcome LbuCas13a-mediated immunity (**Fig. 4D,E**). Therefore, Acrs generated by artificial intelligence can be rapidly deployed as new selectable markers to enable transposition into genomes to engineer phages with expanded anti-defence profiles. These experiments also demonstrate the applicability of the phage Tn*5* Acr transposition system in different genera, including *E. coli* and that phages with hypermodified genomes will be accessible to phage Tn-seq. Together, these features highlight the broad applicability of phage transposon mutagenesis for studying diverse phages and their bacterial host as well as for engineering phages with enhanced therapeutic potential.

## Discussion

While Tn-seq has been instrumental in bacterial genetics, a lack of universal selectable markers hampered its development into a high-throughput method for phages^56–58^. By creating an Acr transposon to counteract a CRISPR-Cas system, we have designed a selection strategy for phage mutagenesis, providing a powerful framework for high-throughput functional genomics and the exploration of gene essentiality in diverse lytic phages. We demonstrated the utility of phage Tn-seq through genome-wide mutagenesis and the delivery of genetic cargo to phages infecting different bacterial genera, including those with either unmodified or hypermodified double-stranded DNA and a nucleus-forming jumbo phage^18^. Furthermore, phage Tn-seq was successfully applied to phages with different DNA packaging and injection mechanisms and can enable the prediction of early injected genome regions.

Across nucleus-forming jumbo phages (*Chimalliviridae*), the fraction of essential genes varies with genome size. In a recent preprint, ~30% of phiKZ genes were reported as essential^59^, whereas our analysis of PCH45 predicted a higher proportion of essential genes (~45%). However, the number of essential genes is similar (~100-110). Furthermore, the PCH45 essentiality data was well supported by the proteins present in mature virions (identified by proteomics), core conserved *Chimillaviridae* genes and essential phiKZ genes^12,59^. Since phiKZ is substantially larger (~370 vs. ~225 genes in PCH45), larger phages appear to encode more accessory or conditionally dispensable genes. This trend is supported by comparisons to smaller (non-jumbo) phages. In phage JS26 (64 kb), ~56% of genes were essential (with high insertion density), while *E. coli* phage T4 (~170 kb) was recently estimated to have ~37% of essential genes^60^. Together, these observations suggest that increasing genome size is associated with a growing pool of non-essential or context-dependent genes.

Phage Tn-seq provides a simple, cost-effective approach for defining gene essentiality in diverse phages and does not require a well-annotated genome. Because it relies on random transposon insertion, precise gene boundaries, start codons, or RBS annotation – requirements for guide design for CRISPRi-based methods^11,61–63^ – are not needed, making Tn-seq suited to non-model or genetically less tractable systems. In contrast to transient knockdown, such as CRISPRi or ASOs^11,12^, Tn-seq generates stable mutations, yielding loss-of-function phenotypes and reducing the likelihood of underestimating essentiality due to incomplete or variable knockdown, which in CRISPRi can be influenced by guide or target RNA secondary structure. Finally, the overall cost of a genome-wide phage Tn-seq experiment, including biological replicates, is approximately half that of comparable CRISPRi-ART screens, while avoiding guide-to-guide variability and the need for multiple guides per gene. In addition, individual phage mutants can easily be picked and purified for use in downstream applications.

There are multiple possible applications and extensions of phage Tn-seq. We used this method to define the essential genes of phage JS26 and a nucleus-forming jumbo phage (PCH45). Since gene essentiality is context-dependent, a strength of Tn-seq is that phage libraries can be applied to different strains or conditions to identify genes influencing host-range or infection. For example, to uncover phage-encoded counter-defence mechanisms, phage transposon libraries could be used to infect different bacteria with distinct immune systems. Indeed, a recent pre-print generated a mariner-based transposon mutagenesis approach of *E. coli* phages and revealed multiple phage counter-defence genes^60^. Stepwise or combinatorial transposon mutagenesis, analogous to Dual Tn-seq in bacteria, could enable genome-wide mapping of functional interactions within phages or between phages and host genes. Our generation of two separate anti-CRISPR selection transposons (Tn*5*-AcrVIA1 and Tn*5*-AIcrVIA3) should allow the generation of phages with double mutations, which could be developed into Dual Tn-seq in the future^64^. The discovery of diverse defence and counter-defence proteins in recent years, as well as the *de novo* design of artificial anti-defence proteins^7,65,66^, offers extensive orthogonal selection and counter-selection markers for further phage Tn-seq development.

We also demonstrated that Tn-seq can rapidly (< one week) facilitate cargo delivery to phage genomes, providing enormous potential for on-demand biotechnological or therapeutic phage development. The addition of counter-defence genes enables phages to evade bacterial immunity, as illustrated in our study. Jumbo phages, such as PCH45, protect their genomes from DNA-targeting CRISPR-Cas and restriction-modification but remain vulnerable to Cas13 (type VI) and type III RNA-targeting systems^21,36,38^. Here, by loading PCH45 with Acrs against Cas13, we broadened the defence evasion by this phage – enabling more robust phages for potential therapeutics. As a further proof-of-concept of cargo delivery, we delivered mEGFP to phages, which will allow fundament studies of phage biology and infection by facilitating microscopy or complementary single-cell techniques. Since transposon mutants that do not adversely affect phage fitness will be preferentially amplified, phages can easily be engineered to carry diverse payloads without prior knowledge of ‘safe harbour’ sites for research or therapeutic purposes.

The phage Tn-seq essentiality maps also reveal large regions of non-essential genes. For example, in PCH45, a large section of 40 genes (gp112-gp151) was non-essential (except gp139). This information could be used to replace large parts of phage genomes with extensive genetic cargo, while still ensuring the phage genome size stays within the head packaging limits. In addition, many of these non-essential genes may be proteins involved in counter-defence or host manipulation and will provide a source of candidate genes for further analysis.

Our study coincides with two recent preprints reporting related transposon sequencing methods for phages^59,60^. Together these three studies provide convincing evidence for the broad applicability of phage Tn-seq methods in different bacteria (*E. coli*, *Pseudomonas* and *Prodigiosinella*) and against diverse phages. While our group employed the Tn5 transposon, the other two groups utilised the mariner transposon. In conclusion, we have developed a fast and efficient genome-wide, high-throughput method with broad utility for investigating phage gene essentiality and for engineering phages with genetic cargo for research or therapeutic applications.

## Methods

### Bacterial growth conditions

All bacterial strains and phages used in this study are listed in **Table S3** and **Table S4**, respectively. *Prodigiosinella confusarubida* sp. ATCC 39006 LacA (formally *Serratia* sp. ATCC 39006) strains were cultured at 30°C and *Escherichia coli* at 37°C in lysogeny broth (LB) with shaking at 180 rpm unless indicated otherwise. For growth on plates, LB supplemented with 1.5% (w/v) agar (LBA) was used. Minimal medium contained 40 mM K_2_HPO_4_, 14.6 mM KH_2_PO_4_, 0.4 mM MgSO_4_, 7.6 mM (NH_4_)_2_SO_4_ and 0.2% (w/v) (w/v) ᴅ-glucose (glucose). When required, media were supplemented with chloramphenicol (Cm, 25 μg/ml), tetracycline (Tc 10 μg/ml or 5 μg/ml), 5-aminolaevulinic acid hydrochloride (Ala, 50 μg/ml), glucose (0.2% w/v), or arabinose (0.2% w/v). Plasmids were introduced into *E. coli* ST18 or DH5α by heat shock or electroporation and transferred into *Prodigiosinella* via conjugation using *E. coli* ST18 as the donor.

### Phage propagation and titration

To generate JS26 and PCH45 phage stocks, a 10 ml overnight culture of *Prodigiosinella* was diluted 1:100 into 50 ml LB and grown to an OD_600_ of 0.2 - 0.3. The culture was then infected with 100 μl of a high-titre phage stock (~10¹⁰ pfu/ml) and incubated overnight (30°C, 180 rpm). Cells were pelleted by centrifugation (3,220 xg, 1 h), and the phage containing supernatant was collected and filtered through a 0.22 μm syringe filter membrane. Phage titres were determined by preparing tenfold serial dilutions in phage buffer (10 mM Tris, pH 7.4; 10 mM MgSO₄, 0.01% w/v gelatin) and spotting 10 μl of each dilution onto 5 ml LB top agar (0.35% w/v or 0.5% w/v) overlays seeded with 150 μl of *Prodigiosinella* overnight culture. Plates were incubated at 30°C overnight, and plaques were counted to calculate titres in pfu/ml.

### Transposon insertion mutagenesis plasmids

All plasmids used or generated in this study are listed in **Table S5**. The phage transposon plasmids were constructed using a combination of Gibson assembly and restriction enzyme–based cloning. The initial transposon backbone, pPF3892, containing an empty transposon, was assembled via Gibson assembly using fragments amplified from plasmids pKRCPN2 (primers PF7472/PF7473) and pPF1618 (primers PF7474/PF7475), together with the synthetic gBlock PF7470. Details of all primers and gBlocks used in this study are provided in **Table S6**. The anti-CRISPR gene *acrVIAI* was subsequently inserted into the transposon by restriction digestion of pPF3892 followed by Gibson assembly with gBlock PF7470, generating plasmid pPF3893. To enable regulated expression of the *tnpA* transposase, plasmid pPF4213 (**Fig. 1B**) was created by replacing the constitutive promoter with the P*_araBAD_* promoter using Gibson assembly of fragments amplified with primer pairs PF7793/PF7888 and PF8074/PF7886 (from pPF3893), together with PF8830/PF8831 (from pPFpBAD30). Plasmid pPF4354 was constructed by fusing the PCH45 gene *uvsX* to *tnpA*, using Gibson assembly of fragments amplified with PF9054/PF9055 (from the PCH45 genome), PF7793/PF9155 and PF9060/PF8598 (from pPF4213), along with the synthetic gBlock PF9066. Finally, plasmid pPF4881 was constructed by Gibson assembly of fragments amplified with primer pairs PF9475/PF7888 and PF9476/PF788 (from pPF4213), together with synthetic gBlock PF9501

### Construction of CRISPR-Cas targeting plasmids

All LseCas13a expression plasmids were derived from pC020 - LseCas13a from *Listeria seeligeri* was a gift from Feng Zhang (Addgene plasmid # 91910; http://n2t.net/addgene:91910; RRID:Addgene_91910)^67^. Plasmid pPF3929 (**Fig. 1C**) was generated by introducing the RP4 oriT into this backbone via Gibson assembly of fragments amplified with PF8098/PF8099 and PF8100/PF8101. New CRISPR spacers were incorporated into this backbone or its derivatives using either primer annealed spacer oligonucleotides and Golden Gate cloning (BsmBI v2) or Gibson assembly when a second spacer cloning site was introduced (**Fig. S1A**). Plasmid pPF4137, containing a spacer targeting the PCH45 major capsid protein (MCP) start codon region, was produced by annealing PF8639/PF8640 and inserting the duplex into BsmBI V2 digested pPF3929. Using this construct, pPF4317 was created by adding a second empty spacer insertion site through Gibson assembly of the annealed primers PF9052/PF9053 into HindIII digested pPF4137. A second targeting spacer (against PCH45 gp033) was added to this background by annealing PF7447/PF8046 and Golden Gate cloning to generate pPF4323. For JS26 targeting constructs, pPF4226 was assembled by inserting the PF8821/PF8822 spacer (targeting JS26 gp82) into BsmBI V2 digested pPF3929. From this plasmid, pPF4364 containing the gp82 spacer and an empty cloning site—was built by Gibson assembly of annealed PF9052/PF9053 into HindIII digested pPF4226. Finally, the dual spacer plasmid pPF4420, encoding spacers targeting JS26 gp82 and gp5, was generated by annealing PF9116/PF9117 and inserting the duplex via Golden Gate cloning into BsmBI V2 digested pPF4364.

### Fluorescent protein plasmid construction

Plasmid pPF4355, a pPF4213 derivative encoding an mCherry2::TnpA fusion, was constructed using Gibson assembly. Fragments were amplified with primer pairs PF9057/PF9058 (from pPF1951), PF7793/PF9156 and PF9060/PF8598 (from pPF4213), and assembled together with the synthetic gBlock PF9065. Plasmid pPF4356, a pPF4213 derivative encoding a UvsX::mCherry2::TnpA fusion along with AcrVIA1, was generated by Gibson assembly of fragments amplified using PF9054/PF9056 (from the PCH45 genome), PF9059/PF9058 (from pPF1951), PF7793/PF9156 and PF9060/PF8598 (from pPF4123), together with gBlock pPF9065. Finally, plasmid pPF4357, a pPF1618 derivative expressing an mCherry2::UvsX fusion, was constructed by amplifying the fusion from pPF4356 using primers PF9094/PF9095 containing AvrII and KpnI overhangs. The resulting PCR product and plasmid pPF1618 were digested with AvrII and KpnI, followed by ligation to generate the final construct.

Plasmid pPF4485, a pPF4213 derivative encoding a UvsX::Tnp fusion, was constructed using restriction digest, An mEGFP insert was amplified with primer pairs pPF9433 and pPF8911 containing restriction sites (from PF1956). Plasmid PF4213 and mEGFP fragment were digested with SacI and NcoI restrictions enzymes and annealed together.

### mCherry fusion confocal microscopy sample preparation

*Prodigiosinella* cells harbouring a mCherry::TnpA fusion (pPF4355), UvsX::mCherry2::TnpA fusion (pPF4356), or mCherry2::UvsX fusion (pPF4357) were grown in triplicate overnight in 5 ml LB supplemented with 5 μg/ml Tc. These cultures were used to seed new 25 ml LB cultures in 125 ml flasks at starting OD_600_ of 0.05 supplemented with 0.2% w/v arabinose for fusion induction and 5 μg/ml Tc for plasmid maintenance. Cultures were grown for ~3.5 h until reaching exponential phase (OD_600_ of 0.3). Phage PCH45 was added at an MOI of 50 and incubated with shaking for 45 min. Following growth, 1.5 ml of each culture was removed to a 1.5 ml microcentrifuge tube and centrifuged at 17,000 ×g to pellet cells. Each pellet was washed 2× with 500 μl minimal medium. Pellets were resuspended in 46 μl minimal medium, and 4 μl of DAPI (final concentration 4 μg/mL) was added to each sample. Samples were incubated protected from light at RT for 5 min, then centrifuged for 30 s at 17,000 ×g. The supernatant was removed and the pellet was washed with 500 μl minimal medium and centrifuged for 30 s at 17,000 ×g. The supernatant was removed and the pellet was resuspended in 50 μl of minimal medium. To resuspend samples, 15 μl aliquots of each sample were mixed with 15 μl of molten 1.2% (w/v) agarose (in minimal medium) before being sealed onto microscope slides with a coverslip.

### mEGFP fusion confocal microscopy sample preparation

*Prodigiosinella* cells harbouring a PCH45 LseCas13a targeting plasmid (pPF4324), or JS26 LseCas13a targeting plasmid (pPF4420) were grown for confocal microscopy as stated above. Phage PCH45 and JS26 harbouring Tn*5*-mEGFP-AcrVIA1 were picked as single plaques following enrichment, into 100 µl of phage buffer, 90 µl of which was inoculated into 5 ml of cultures containing their respective targeting plasmid (pPF4324 and pPF4420) at an OD_600_ of 0.3. Phage enrichments were grown for ~5hrs then syringe-filtered (0.22 µm). *Prodigiosinella* cells were then infected with 1 ml of filtered phage lysates (1 ml) for 45 min. Infected cells were then collected, stained, and mounted for imaging as described above.

### Confocal microscopy and image analysis

Samples were imaged and processed as previously described^38^. Briefly, images were acquired using a CFI Plan APO Lambda ×100 1.45 numerical aperture oil objective (Nikon Corporation) on the multimodal imaging platform Dragonfly v.505 (Oxford Instruments) equipped with 405, 488, 561 and 637 nm lasers built on a Nikon Ti2-E microscope body with Perfect Focus System (Nikon Corporation). Data were collected in Spinning Disk (40 μm pinhole, 133 nm spacing) mode on the iXon888 EMCCD camera with x2 optical magnification using the Fusion Studio Software v.1.4 (Andor Oxford Instruments). Z stacks were collected with 0.1 μm increments on the z-axis using an Applied Scientific Instrumentation stage with 500 μm piezo z drive. Images were visualized and cropped using Fiji (ImageJ2) v.2.16.0 and processed using the Spinning Disk Deconvolution Wizard of Huygens Essential (Scientific Volume Imaging). Final composite images and fluorescence plot data were generated using Fiji and graphed using Prism v.10.5.0 (GraphPad).

### Efficiency of plaquing assays

To determine targeting efficiency of *Prodigiosinella* Type VI CRISPR-Cas targeting strains containing LseCas13a expression plasmids with either one or two gRNA inserts, efficiency of plaquing (EOP) assays were performed. Overnight cultures of *Prodigiosinella* Type VI CRISPR-Cas targeting strains were grown in LB supplemented with Tc at 30°C with shaking (120rpm). A molten LBA overlay (5 ml, 0.35% w/v) previously seeded with 150 µl of the respective bacterial culture was poured onto LBA plates. Serial tenfold dilutions of transposon phage lysate were spotted (5 µl) onto the agar overlay and plates were incubated overnight at 30°C. The EOP was calculated as a ratio of PFU/ml produced on the respective CRISPR targeted strain and the PFU/ml on the untargeted control. n=3 independent biological replicates.

### Phage transposon mutagenesis

An overview of the transposon mutagenesis methodology is provided in Figure 1A and Figure S1D. The transposon saturated mutant phage pools (n=3 independent biological replicates) were generated by first inoculating *Prodigiosinella* harbouring the either the transposon plasmid pPF4213 or pPF4354 into 50 ml of Luria Broth (LB) supplemented with 0.2% w/v glucose and 5 mg/ml Tc. Cultures were grown at 30°C with shaking (160 rpm) until reaching an OD_600_ ≥ 2, then harvested by centrifugation (4,000 × g, 10 min), washed once with 25 ml LB, pelleted again, and resuspended in 25 ml LB. The culture was used to inoculate 500 ml of LB and Tc at a starting OD_600_ of 0.1. The culture was grown for 30 min at 30°C, shaken, before 0.2% w/v arabinose was added for transposase induction. The culture was then returned to the incubator and grown until an OD_600_ of 0.3 at which phage were inoculated with a multiplicity of infection (MOI) of 1. Infecting cultures were then incubated for 18 h, phage were collected through centrifugation and filtering, and titred as above.

### Enrichment of mutagenised phage

The transposon mutagenised phages were enriched through negative selection against the non-transposon carrying phage. *P. confusarubida* carrying a LseCas13a targeting plasmid was inoculated in 50 ml of LB supplemented with Cm incubated at 30°C shaken at 160 rpm overnight. The host was grown in 50 ml of LB supplemented with Cm incubated at 30°C shaken at 160 rpm overnight. The overnight culture was then used to inoculate 500 ml of LB supplemented with Cm at an OD_600_ of 0.05. The culture was then grown to an OD_600_ of 0.3, transposon phage culture was then inoculated at an MOI=1 and incubated for 18 h.

### PCR of phage plaques for transposon insertions

To test phage plaques for successful transposon insertions phage plaques were picked from agar overlays using the same method as for phage titration, except serial dilutions of 100 µl of phage stock was added to the overlay before it was poured. Plates containing spaced individual plaques were then used for individual plaque isolation. Plaques were picked and resuspended in either 100 µl of phage buffer (and 5 µl added to PCR master mix) or in PCR master mix directly with screening primers PF7658 and PF8715 (AcrVIA1), PF7793 and 8715 (AIcrVIA3), or PF9057 and PF9058 (mEGFP) and run on an agarose gel.

### Phage DNA extraction

Phage genomic DNA was extracted from high-titre lysates following ultracentrifugation, nuclease treatment, and organic extraction. Briefly, clarified lysates were transferred to Ti-70 ultracentrifuge tubes and phage particles were pelleted by ultracentrifugation at 40,000 rpm for 30 min. Pellets were resuspended in 500 μl nuclease-free water (with softening at 4°C if necessary), and residual bacterial or plasmid nucleic acids were removed by treating 5 ml lysate with RNase A (5 μl), DNase I (>2 U; 5 μl of 1 mg ml⁻¹), and 10× DNase buffer (500 μl) at 37°C for 30 min. Nucleases were inactivated by adding 400 μl of 0.5 M EDTA and incubating at 65°C for 10 min, after which phage capsids were lysed with proteinase K (62.5 μl of 10 mg ml⁻¹) at 45°C for 15 min. DNA was purified by phenol–chloroform extraction: samples were mixed with an equal volume of phenol:chloroform:isoamyl alcohol, centrifuged (3,000 × g, 10 min), and the aqueous phase recovered, followed by a second extraction with chloroform alone. DNA was precipitated by adding 0.1 volumes of 3 M sodium acetate (pH 5.0) and 2.5 volumes of ice-cold ethanol, incubating at −20°C for ≥1 h, and pelleting at 14,000 × g (4°C). Pellets were washed with 70% ethanol, briefly air-dried, and resuspended in 400 μl nuclease-free water, with overnight incubation at 4°C if required for complete dissolution. DNA quantity and purity were assessed using a NanoDrop spectrophotometer or Qubit fluorometer.

### Library preparation and sequencing

Sequencing libraries were prepared using the NEBNext Ultra II FS DNA Library Prep Kit for Illumina following established protocols^18^. In brief, library enrichment was performed in two PCR steps. The first amplification used primer PF3140, which anneals to the Illumina adaptor, together with biotinylated primer PF8656 targeting the transposon; all primers are listed in Supplementary Table S3. Biotin-labelled amplicons were isolated using Dynabeads M-270 Streptavidin (Invitrogen) according to the manufacturer’s guidelines, and the bead-bound DNA served as template for a second PCR using a nested transposon primer (PF8657) and a unique index primer (NEBNext Multiplex Oligos). Library size distribution and integrity were evaluated using an Agilent 2100 Bioanalyzer with the High Sensitivity DNA Kit. Libraries were quantified by Qubit fluorimetry (dsDNA HS Kit, Thermo Fisher Scientific), diluted to 10 nM based on Qubit concentration and average fragment length, and pooled with 10% PhiX control. The final pool was loaded at 1.5 pM (low-diversity loading) and sequenced on an Illumina Miseq V2 standard kit. Single-end 75-cycle reads were generated with custom sequencing primer PF2926 and the standard Illumina Read 1 primer PF3441 (for PhiX). Sequencing with PF2926 produces a 12-nt transposon tag, enabling the identification of transposon–genome junction reads.

### Tn-seq data analysis

Samples were de-multiplexed based on index sequence using standard Illumina software. FASTQ files were trimmed using Trimmomatic v.0.39 and mapped to reference sequences using the Bio-Tradis pipeline v.1.4.5^19^. The following parameters were used to run the bacteria_tradis script, which identifies the transposon tag in FASTQ files, then maps reads to the reference genome: ‘–smalt –smalt_k 10 –smalt_s 1 –smalt_y 0.92 -mm 2 -v -f filelist.txt -t TATAAGAGACAG -r PCH45/JS26.fasta’, where -smalt specifies the SMALT read mapping tool (https://www.sanger.ac.uk/tool/smalt/), -smalt_k specifies kmer size for reference fasta, -smalt_s specifies step size for kmers, -smalt_y specifies minimum percent identity of identical bases between read/reference, -mm specifies mismatches allowed in transposon tag sequence, -smalt_r specifies that reads that map equally in multiple places are discarded, -t specifies the transposon tag sequence, -r specifies the reference sequence, and -f indicates the list of FASTQ files to be processed. In summary, only reads containing the Tn tag (minimum 10/12 identity) and 92% identity to the reference sequence were considered for further analysis. Plot files (.insert_site_plot), which tabulate the number of reads at each nucleotide position in the reference sequence (plus and minus strands), were generated by the bacteria_tradis script. Reads mapping to the 3’ end of genes (10% of the feature length) were trimmed using ‘tradis_gene_insert_sites - trim3 0.1 PCH45/JS26.embl *.insert_site_plot.gz’ which were then analysed with the tradis_essentiality.R script for outputs of essential and ambiguous genes. Summaries of the mapping data are provided in Tables S1 and S2 for phages JS26 and PCH45, respectively. Summaries of per-gene insertion data (pre- and post-trimming) are provided in Extended Data Tables 1 and 2 for phages JS26 and PCH45, respectively. Data was further processed and visualised using R v.4.5.1 RStudio (2025 09.1 build 401).

### Phage JS26 RNA-seq data analysis

Previous RNA-sequencing analysis of a phage JS26 infection time course (5, 20, and 40 min post-infection) in *P. confusarubida* assigned each JS26 gene to different temporal expression clusters (early; gp01-14, middle; gp15-gp45, late; gp46-gp74, or middle-late; gp75-gp85)^26^. To visualise expression data (as seen in Figure 2B), we calculated the mean of normalised count data (n=3) for each gene during its assigned temporal cluster (early; 5 min, middle; 20 min, late; 40 min, and middle-late; 40 min post-infection). The mean was then transformed using the following formula: log10((mean normalised counts / gene length)+1).

### Phage JS26 genome end analysis

Previously-published whole genome DNA-sequencing reads of phage JS26^26^ were mapped to the circularised JS26 reference sequence using Geneious Prime v.21.0.04. Read coverage was manually inspected, with a drop in coverage and reads spanning the ends detected.

### Phage PCH45 structural proteomics

Phage PCH45 virions were purified via sucrose cushion gradient ultracentrifugation as previously described^26^. Proteomic analysis of the phage structural proteins was performed as previously described^26,68^. Briefly, 75 µl of purified phage sample was mixed with 20 µl of 1/10 (v/v) β-mercaptoethanol 4x SDS-buffer and incubated for 5 min at 95⁰C. The sample was separated by electrophoresis in a 12% w/v SDS-polyacrylamide gel for 1 h at 150 V. Page ladder (Thermo Fisher Scientific) was used as a reference. The gel was fixed for 2 h in a 10% v/v acetic acid, 40% v/v ethanol. Finally, the gel was stained overnight in 4 parts of Colloidal Coomasie Blue stain (0.1% G250 w/v, 10% w/v ammonium sulphate and 2% v/v orto-phospholic acid) and 1 part methanol. The gel lane was subjected to in-gel digestion with trypsin and analysed by protein identification by liquid chromatography-coupled tandem mass spectrometry (LC-MS/MS). Peptide reconstruction was performed with an Ultimate 3000 nano-flow uHPLC-System (Dionex Co, Thermo Fisher Scientific) in-line coupled to the nanospray source of an LTQ-Orbitrap XL mass spectrometer (Thermo Scientific; Waltham, USA). Raw spectra were processed through the Proteome Discoverer software (Thermo Fisher Scientific) using default settings to generate peak lists. Peak lists were then searched against a combined amino acid sequence database containing all PCH45 sequence entries (GenBank accession number MN334766.1, 225 entries) integrated into the full SwissProt/UniProt sequence database using the Sequest HT (Thermo Fisher Scientific), Mascot (www.matrixscience.com) and MS Amanda search engines (**Extended Data Table 3)**.

### Hypothetical protein functional assignments

Putative functions were assigned to hypothetical genes through protein sequence and structure analyses. Protein sequences were initially searched against public databases using NCBI BLAST+ (v.2.17.0) to identify homologous proteins based on sequence similarity. Remote homology detection was performed with HHpred via the HH-suite v.3.3.0 backend to detect relationships through profile HMM comparisons^69^. Predicted protein structures were input into AlphaFold protein structure database (v.6) and used to identify structural homologs and infer potential functions^70^. Additionally, structural similarity searches were performed using the Foldseek web server (Release 10-941cd33) to compare predicted structures against known protein structures^71^.

## Acknowledgements

This research was supported by a James Cook Research Fellowship (RSNZ, Te Apārangi) to PCF and by research funds from the Otago Medical School Foundation Trust or Otago Faculty of Biomedical Sciences Dean’s Fund Awards (LMS and PCF). NK was supported by University of Otago Doctoral scholarship. LMS was supported by the Marsden Fund and a L’Oréal-UNESCO For Women in Science Fellowship (The L’Oréal Groupe Australia). MF was supported by a Feodor Lynen Research Fellowship from the Alexander von Humboldt Stiftung. We thank Rob Wooley and Rose Smither of the Otago Micro and Nano Imaging (OMNI) facility for assistance with Confocal Microscopy, Monika Zavodna of the Otago Genomics Facility (OGF) for genome sequencing, Torsten Kleffmann of the Centre for Protein Research (CPR) for proteomics and staff of the Genetic Analysis Service (GAS) for Sanger sequencing. Thanks to Lucia Malone for the preparation of phage PCH45 samples for proteomic analyses.

## Author Contributions

Conceptualisation, PCF; Formal analysis, NK, MF, LMS, PCF; Funding acquisition, LMS, PCF; Investigation, NK, MF, LMS; Project administration, LMS, PCF; Supervision, LMS, PCF; Visualisation, NK, MF, LMS; Writing – original draft, all authors; Writing – review & editing, all authors.

## Competing Interests

The authors declare no competing interests.

## Additional Information

### Supplementary information

#### Extended data

**Extended Data Table 1. Phage JS26 gene essentiality, proteomics and RNA-sequencing data summary.** For transposon insertion sequencing data, read counts (read_count), unique insertions (insertion_count), insertion index (insertion_index; insertion_counts / gene_length) and essentiality predictions (essentiality; Essential, Non-essential or Ambiguous) are indicated for each gene (gene_name), for both raw insertion mapping (_raw) and trimmed (_trimmed), where reads/insertions mapping to 10% of the gene feature (from the 3’ end) were trimmed. The presence of a gene product in the mature virion (structural_proteomics) is indicated by Yes/No. Each gene was assigned to an expression block (gene_expression_timing; Early, Middle, Late, Middle-Late) based on RNA-sequencing data^20^. The mean gene expression across (*n*=3) replicates are shown for each time point (Early; 5 min, Middle; 20 min, Late; 40 min-post infection).

**Extended Data Table 2. Phage PCH45 gene essentiality, proteomics and core genome predictions data summary.** For transposon insertion sequencing data, read counts (read_count), unique insertions (insertion_count), insertion index (insertion_index; insertion_counts / gene_length) essentiality predictions (essentiality; Essential, Non-essential or Ambiguous) are indicated for each gene (gene_name), for both raw insertion mapping (_raw) and trimmed (_trimmed), where reads/insertions mapping to 10% of the gene feature (from the 3’ end) were trimmed. The presence of a gene product in the mature virion (structural_proteomics) is indicated by Yes/No. Genes were assigned to the *Chimalliviridae* family predictions^45^ including the nucleus-forming ‘core’ genome (core_genome_predictions) and the subset of core genes unique to Chimallin-encoding phages (chimallin_genome_predictions). The core genome number (Cg) and putative function is indicated (core_genome_homolog).

**Extended Data Table 3. Phage PCH45 proteomics data.** Summary of mass spectrometry results from the analysis of purified phage PCH45 particles. Per protein detected by mass spectrometry the following columns of information are provided: 1) gene name, 2) gene description, 3) essentiality (trimmed 10% 3’ of gene), 4) jumbo phage ‘core’ genes or 5) core genes unique to *Chimallin*-encoding phages genome predictions^45^, 6) core genome number and annotation^45^, 7) Unique peptides identified per protein, 8) The number of acquired peptide spectra per protein (PSMs-peptide spectral matches), resulting in the identified 9) sequence coverage (coverage (%)), 10) Protein score calculated as the negative logarithms of the Posterior Error Probability values of PSMs and 11) Score calculated by search engine Mascot.

**Correspondence and requests for materials** should be addressed to

## Notes

### Competing Interest Statement

The authors have declared no competing interest.

